# Quantification of influenza virus mini viral RNA dynamics using Cas13

**DOI:** 10.1101/2023.11.03.565460

**Authors:** Caitlin H. Lamb, Emmanuelle M. Pitré, Elizaveta Elshina, Charlotte V. Rigby, Karishma Bisht, Michael S. Oade, Hamid Jalal, Cameron Myhrvold, Aartjan J.W. te Velthuis

**Affiliations:** Department of Molecular Biology, Princeton University, Princeton, NJ 08544; University of Cambridge, Department of Pathology, Addenbrooke’s Hospital, Cambridge CB2 2QQ, United Kingdom; Public Health England, Addenbrooke’s Hospital, Cambridge CB2 2QQ, United Kingdom; Department of Chemical and Biological Engineering, Princeton University, Princeton, NJ 08544; Omenn-Darling Bioengineering Institute, Princeton University, Princeton, NJ 08544; Department of Chemistry, Princeton University, Princeton, NJ 08544

**Keywords:** influenza virus, mini viral RNA, RNA polymerase, Cas13, CRISPR, innate immune, RIG-I, t-loop

## Abstract

Influenza A virus RNA synthesis produces full-length and aberrant RNA molecules, which include defective viral genomes (DVG) and mini viral RNAs (mvRNA). Sequencing approaches have shown that aberrant RNA species may be present during infection, and that they can vary in size, segment origin, and sequence. Moreover, a subset of aberrant RNA molecules can bind and activate host pathogen receptor retinoic acid-inducible gene I (RIG-I), leading to innate immune signaling and the expression of type I and III interferons. Understanding the kinetics and distribution of these immunostimulatory aberrant RNA sequences is important for understanding their function in IAV infection. Here, we use an amplification-free LbuCas13a-based detection method to quantify mvRNA amplification dynamics and subcellular distributions. We show that our assay can quantify the copy numbers of specific mvRNA sequences in infected tissue culture cells, ferret upper and lower respiratory tract tissue infected with two different pandemic H1N1 IAV strains, or clinical nasopharyngeal swab extracts of hospitalized patients infected with seasonal H1N1 or H3N2 strains. In addition, we find dynamic differences between immunostimulatory and non-immunostimulatory mvRNAs, as well as among mvRNAs derived from different segments, during IAV infection. Overall, our results reveal a hitherto hidden diversity in the behavior of IAV mvRNAs and suggest that individual aberrant RNAs are not produced stochastically.

## Introduction

Influenza A viruses (IAV) cause a mild to severe respiratory disease in humans, depending on the viral strain and activation of the innate immune response. Upon infection, IAV releases eight segments of negative-sense, single-stranded RNA that are organized into viral nucleoprotein (vRNP) complexes. Each vRNP complex consists of a viral RNA (vRNA) that is bound by a helical coil of nucleoproteins (NP) and a copy of the RNA-dependent viral RNA polymerase (*1*, *2*) (Fig. 1A). During replication, the IAV RNA polymerase can produce a wide variety of aberrant RNA products including defective viral genomes (DVG) and mini viral RNAs (mvRNA), which contain the conserved termini of the viral genome segments but lack internal sequences (*3–7*) (Fig. 1A). It is currently assumed that the internal deletions are the result of an intramolecular copy-choice recombination event that involves pausing of the RNA polymerase at an unknown signal and realignment of the nascent RNA to a complementary sequence downstream. Deletion of an internal sequence, results in the formation of a unique junction sequence relative to the full-length viral genome (Fig. 1B).

**Figure 1.**
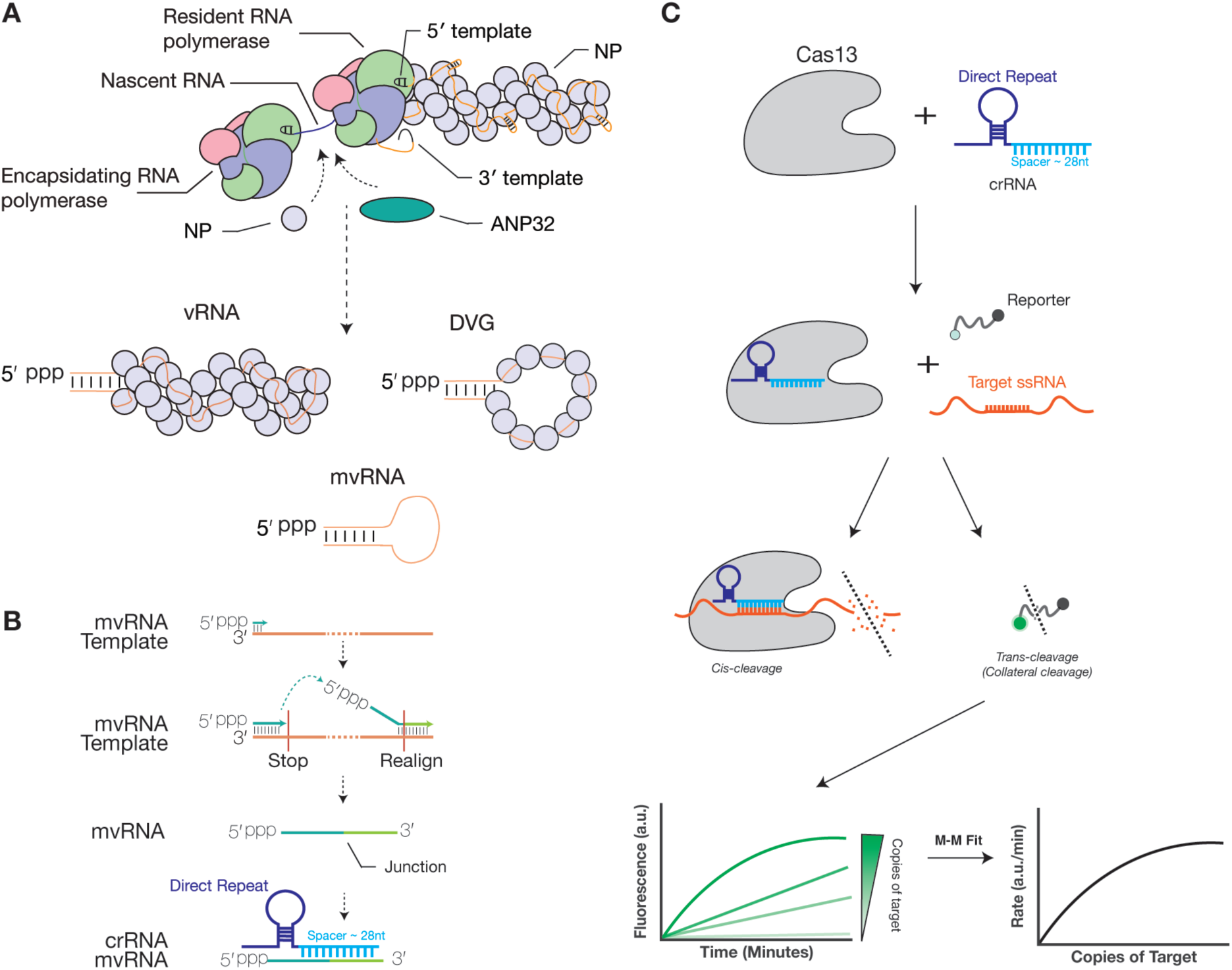
Schematic of influenza A virus RNA replication products and detection mvRNAs using Cas13. **A**) The IAV RNA polymerase forms a dimer during replication. This dimer is stabilized by host factor ANP32A/B and able to recruit viral nucleoprotein (NP) to encapsidate nascent viral RNA. In addition to producing full length RNA products, IAV replication also produces mini viral RNAs (mvRNA) and defective viral genomes (DVG). **B**) During IAV replication mvRNAs can be generated through a copy-choice recombination mechanism that deletes internal genome segment sequences. crRNAs were designed to target the unique junction sequences, thereby distinguishing mvRNAs from full length viral genome segments and other mvRNAs. **C**) Schematic of RNA detection using Cas13.

Some of these aberrant RNAs have been shown to bind and activate retinoic-acid inducible gene I (RIG-I), leading to downstream innate immune signaling (*5*, *8–11*). The production of aberrant RNAs plays a still poorly understood role in viral pathogenicity and the outcome of influenza virus infection (*4*, *5*, *10*, *12*). In the case of mvRNAs, their abundance was associated with the appearance of disease markers in mouse and ferret infections with highly pathogenic IAV strains (*5*, *10*). However, not all aberrant RNA molecules are able to induce an innate immune response. In IAV, as well as other negative sense RNA viruses, sequence-specific preferences for RIG-I binding and activation have been observed (*6*, *9*, *13*, *14*). For IAV, we recently showed that mvRNAs containing a template loop (t-loop), a transient RNA structure that can affect RNA polymerase processivity, are more potent inducers of the innate immune response than mvRNAs without a t-loop in cell culture (*9*), although the sensitivity to t-loops is IAV RNA polymerase-dependent (*15*). Given the large variety of aberrant RNA sequences produced during IAV infection and the specific effects of different sequences, it is becoming more important to carefully study the kinetics and impact of individual aberrant RNA sequences during infection.

Current tools to detect and quantify aberrant RNAs, such as mvRNAs, include primer extension, reverse transcription quantitative PCR (RT-qPCR), and next generation sequencing (*16*). To complement the above assays and perform aberrant viral RNA quantification without RT and PCR-based amplification, we explored the use of Cas13, an RNA-guided nuclease, for the direct, amplification-free detection and quantification of aberrant IAV RNAs (Fig. 1C). Cas13 uses a CRISPR RNA (crRNA) that consists of a direct repeat region, which is specific to the Cas13 ortholog, and a spacer region, which is designed to be complementary to the target RNA (*17*, *18*). When CRISPR-Cas13 binds to the target RNA, the nuclease activity of Cas13 is activated. Cas13 can subsequently cleave the target, a process that is called *cis*-cleavage, as well as any nearby single-stranded RNA molecules, a process called trans-cleavage or collateral cleavage (*19*). The trans-activity can be measured with a reporter RNA molecule containing a fluorophore and a quencher (Fig. 1B), making Cas13 suitable for a wide-range of applications, including the detection of genomic viral RNA (*20–25*). Additionally, the measured fluorescent signal is proportional to the amount of target RNA present in a sample and the exact number of target RNA molecules present in a reaction can be calculated using a standard curve specific for the target RNA (Fig. 1B) (*26*).

Here, we used amplification-free, a Cas13-based detection method to quantify the expression levels for 10 mvRNAs derived from five different IAV segments. We chose to perform this comparison on mvRNAs, which, because their short length (approx. 56-125 nt), facilitated chemical RNA synthesis and thus the possibility to use a well-defined input that could be used to validate and compare assays. We designed our crRNA such that it targeted the unique junction sequence formed upon mvRNA production (Fig. 1B, C) to differentiate mvRNAs from genomic RNA and different mvRNAs from each other. We tested our assays using total RNA from cell culture, animal tissue, or clinical nasopharyngeal swabs, and were able to detect mvRNAs in all sample types. Moreover, we observe kinetic and subcellular distribution differences between immunostimulatory and non-immunostimulatory mvRNAs, as well as mvRNAs derived from different segments. With one exemption, these differences correlate with a previously identified RNA structure that modulates IAV RNA polymerase activity, a so-called template loop (t-loop). These results provide new insight into the diversity and behavior of small, aberrant viral RNAs that are produced during IAV infection in cell culture, ferrets, and humans.

## Results

### Amplification of mvRNAs by RT and PCR introduces errors

Different detection methods can be used to quantify IAV aberrant RNA species, but their errors have not previously been compared directly for mvRNAs. To gain insight into the errors generated by these methods, we compared the ability of RT and PCR to quantify known, chemically synthesized mvRNA sequences. Using two mvRNAs of the same length, but a different internal sequence, we observed a lack of consistency in the ability of three different RT and PCR enzyme combinations to produce a single amplification product (Fig. S1A). Additionally, the different RT and PCR enzyme combinations amplified the mvRNAs to different levels even though a fixed amount of the synthetic mvRNAs was provided as input (Fig. S1A, S1B). Thus, while primer extension, PCR, and next generation sequencing are powerful methods to visualize or discover aberrant IAV RNAs, their use as quantification methods appears limited by the enzymes used for the conversion or amplification of different mvRNA sequences.

### LbuCas13a can be used to detect and quantify mvRNAs without amplification

Cas13 can be used to detect SARS-CoV-2 RNA without (isothermal) amplification and T7 transcription steps when the genome is targeted by multiple guides. As only one unique junction is available in mvRNA sequences (Fig. 1B, C), the previously described approach is not feasible for IAV aberrant RNA molecules. However, Cas13 orthologs have variable amounts of *trans*-cleavage activity (*19*) and this activity can be used to generate additional signal. To determine if Cas13 could be used to detect mvRNAs junctions without the previously described amplification and T7 transcription steps, we compared the ability of two Cas13 orthologs, LbuCas13a and LwaCas13a. First, we compared the activity of the two enzymes in their ability to detect human 5S rRNA in a dilution series of total RNA extracted from HEK293T cells. We observed that LbuCas13a was able to detect 5S rRNA with as little as 0.025 ng of total RNA input, whereas LwaCas13 only produced a signal above background when incubated with 10 ng of total RNA input (Fig. 2A).

**Figure 2.**
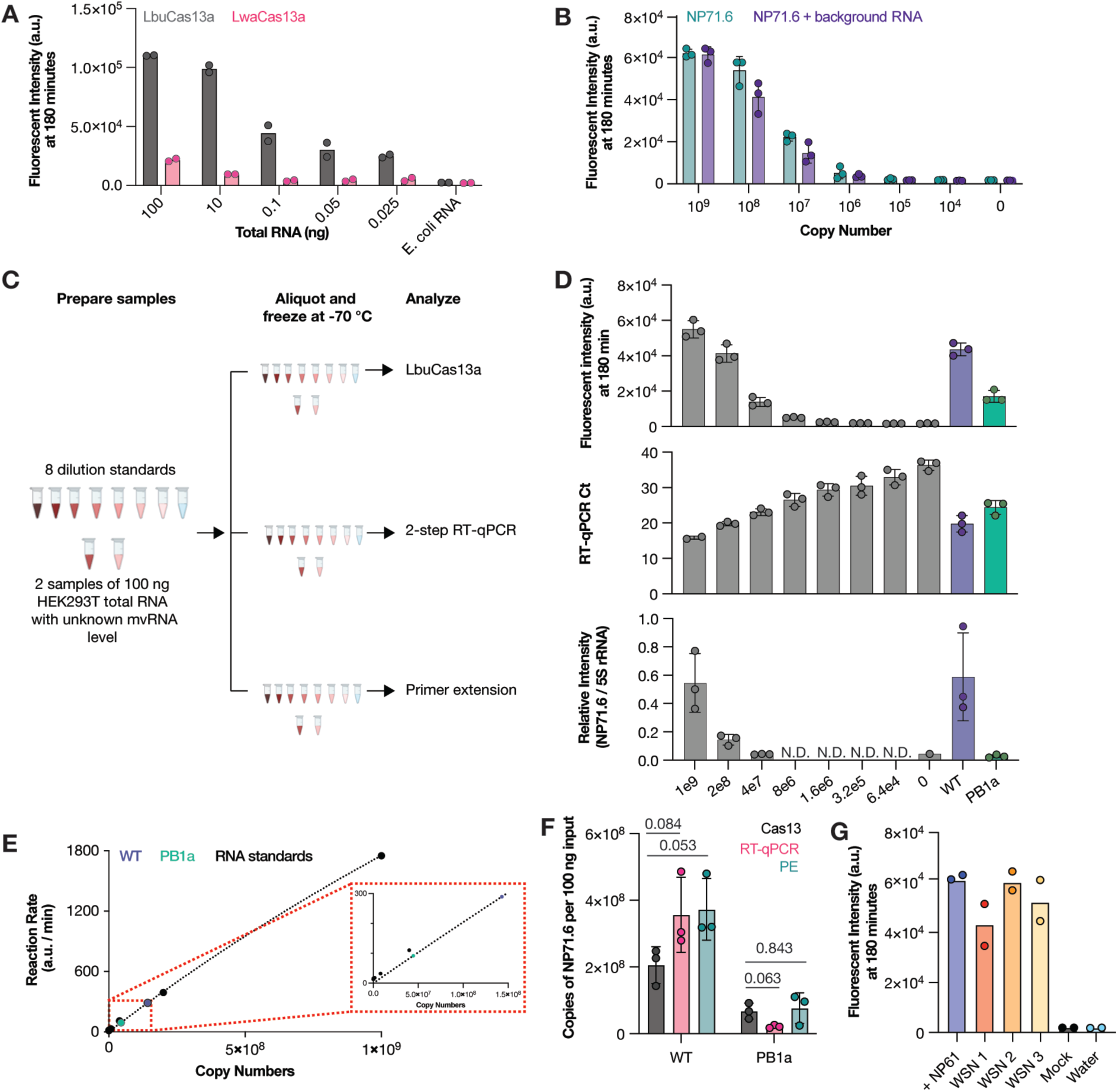
Detection of mvRNAs using LbuCas13a. **A**) Detection of 5S rRNA using LbuCas13a or LwaCas13a. Data points indicate two biological repeats. **B**) Detection synthetic mvRNA NP71.6 diluted in water or HEK293T total RNA using LbuCas13a. Data points indicate three independent repeats. **C**) Schematic of RNA sample preparation for mvRNA detection. **D**) Detection of chemically synthesized (gray bars) or transfected mvRNA NP71.6 in the presence of wildtype or mutant IAV RNA polymerase (purple or green bar, respectively) using LbuCas13a (top), TaqMan-based RT-qPCR (middle) or primer extension (bottom). Data points indicate three independent repeats. For raw primer extension data see Fig. S2. **E**) Michaelis-Menten fits to the maximum rate of fluorescence as a function of synthetic mvRNA NP71.6 copy number. **F**) Copy number of mvRNA NP71.6 in transfection samples. Data points indicate three biological repeats. Statistical comparisons are based on non-parametric t-test. **G**) Copy number of synthetic mvRNA NP-61 or mvRNA NP-61 in WSN infections of A549 cells. Data points indicate two technical repeats. In all graphs, error bars indicated standard deviation.

We next investigated whether LbuCas13a could detect a previously described abundant control mvRNA derived on segment 5 (NP71.6). To detect NP71.6, we designed a crRNA with a spacer sequence complementary to the unique junction sequence relative to the full-length segment 5 vRNA. We subsequently diluted chemically synthesized NP71.6 in water or 100 ng of total HEK293T RNA and found that we were able to detect the mvRNA with a limit of detection <10^6^ copies (Table S2), irrespective of the background used (Fig. 2B). No fluorescent signal was observed in the absence of synthetic NP71.6 (Fig. 2B). We subsequently explored whether we could use LbuCas13a to quantify unknown amounts of NP71.6 in total RNA extracted from HEK293T cells expressing the IAV mini-genome system (*16*). In this assay, plasmids expressing the three IAV RNA polymerase subunits via an RNA polymerase II (Pol II) promoter and the NP71.6 mvRNA via Pol I promoter were transfected into HEK293T cells. Total RNA was extracted 48-hours post transfection. To investigate our ability to determine the mvRNA level in HEK293T cells, we first made a 5-fold dilution series of chemically synthesized NP71.6 in total RNA from untransfected HEK293T cells. This dilution series was our standard curve (Fig. 2C; S2). Next, we calculated the maximum slope of the fluorescent Cas13 signal for each point of the standard curve, plotted the maximum slope against the known synthetic mvRNA concentrations, and fitted the resulting distribution to the Michaelis-Menten equation (Fig. 2D). Next, the Cas13 signal of the transfected HEK293T RNA was determined and the fitted standard curve data used to convert the fluorescent signal of the transfected HEK293T cells into mvRNA copy numbers (Fig. 2D, E).

In Fig. S1, we showed that RT-PCR detection methods can introduce mvRNA amplification biases. To reduce this bias and detect mvRNAs using RT-qPCR in a sequence-specific manner, similar to our Cas13 assay, we introduced a TaqMan probe that was specific for the junction in the NP71.6 mvRNA. When we next compared the quantification obtained using LbuCas13a to the results obtained using primer extension or a two-step TaqMan-based RT-qPCR, we observed indeed that our CRISPR-based detection method yielded no significantly different results (Fig. 2D, E). A similar result was obtained when we expressed the NP71.6 mvRNA in the presence of an inactive IAV RNA polymerase to measure RNA polymerase I (PolI)-derived input levels (Fig. 2E). Consequently, for a single, well-defined IAV mvRNA the RT-qPCR and Cas13 assays yield comparable results. However, in infection experiments, the mvRNA diversity is relatively large, making it more likely that sequence-based biases arise when amplification steps are used for mvRNA detection. We therefore decided to continue our analysis with the amplification-free Cas13 assay only.

We subsequently investigated whether LbuCas13a was able to detect a specific mvRNA sequence during IAV infection. To increase the likelihood of detection, we designed a crRNA complementary to a 61 nt-long, highly abundant mvRNA derived from segment 5 (NP-61), which we had previously identified in ferret lung tissues infected with A/Brevig Mission/1/1918 (H1N1) and A549 cells infected with A/WSN/1933 (H1N1) (abbreviated as WSN) (*5*, *9*). As shown in Fig. 2F, LbuCas13a successfully detected synthetic NP-61 as well as NP-61 expressed during WSN infection of A549 cells in total RNA. No signal was detected in our water and mock infection controls (Fig. 2F). Taken together, the above results indicate that LbuCas13 can detect a specific mvRNA sequence in well-defined samples similar to primer extension and RT-qPCR, and that it can detect a specific mvRNA sequence in a complex RNA sample without sequence-specific biases that are potentially associated with RT and PCR steps, as shown in Fig. S1A and S1B.

### mvRNAs can be detected from 3 hours post infection and accumulate over time

We previously identified many hundreds of different mvRNA sequences using next generation sequencing (*5*, *9*). We and others have proposed that DVGs and mvRNAs are generated through an error-prone mechanism, suggesting that their expression is stochastic. To investigate this hypothesis, we next sought to quantify different mvRNAs produced over the course of IAV infections and study their kinetics. We therefore focused on four sets of two mvRNA sequences derived from IAV segment 2, 3, 4 and 8 (encoding viral proteins PB1, PA, HA, and NS), covering the spectrum of genome segment sizes, segment-specific kinetic differences (*27*), and segment-specific unique mvRNA counts (*9*). Moreover, for each segment, we selected one mvRNA with a t-loop, making it likely that their amplification was reduced and ability to activate the innate immune response increased, and one mvRNA without a t-loop, increasing the likelihood that these mvRNAs were amplified quickly and not able to activate the innate immune response (Fig. 3A, colored red or blue, respectively). We cloned these eight mvRNAs into PolI-driven expression plasmids and confirmed their ability or inability to activate innate immune signaling by transfecting the plasmids in HEK293T cells alongside constructs expressing the IAV RNA polymerase subunits, a *Renilla* luciferase transfection control, and a luciferase based IFNβ reporter. Measurement of the luciferase signal showed that two mvRNAs, derived from segment 1 and 2 (PB1-66 and PA-66, respectively) were strong inducers of IFN-β promoter activity, while the other six mvRNAs were relatively weak or non-inducers (i.e., PB1-64, PA-60, HA-61, HA-64, NS-56 and NS-80) (Fig. 3A).

**Figure 3.**
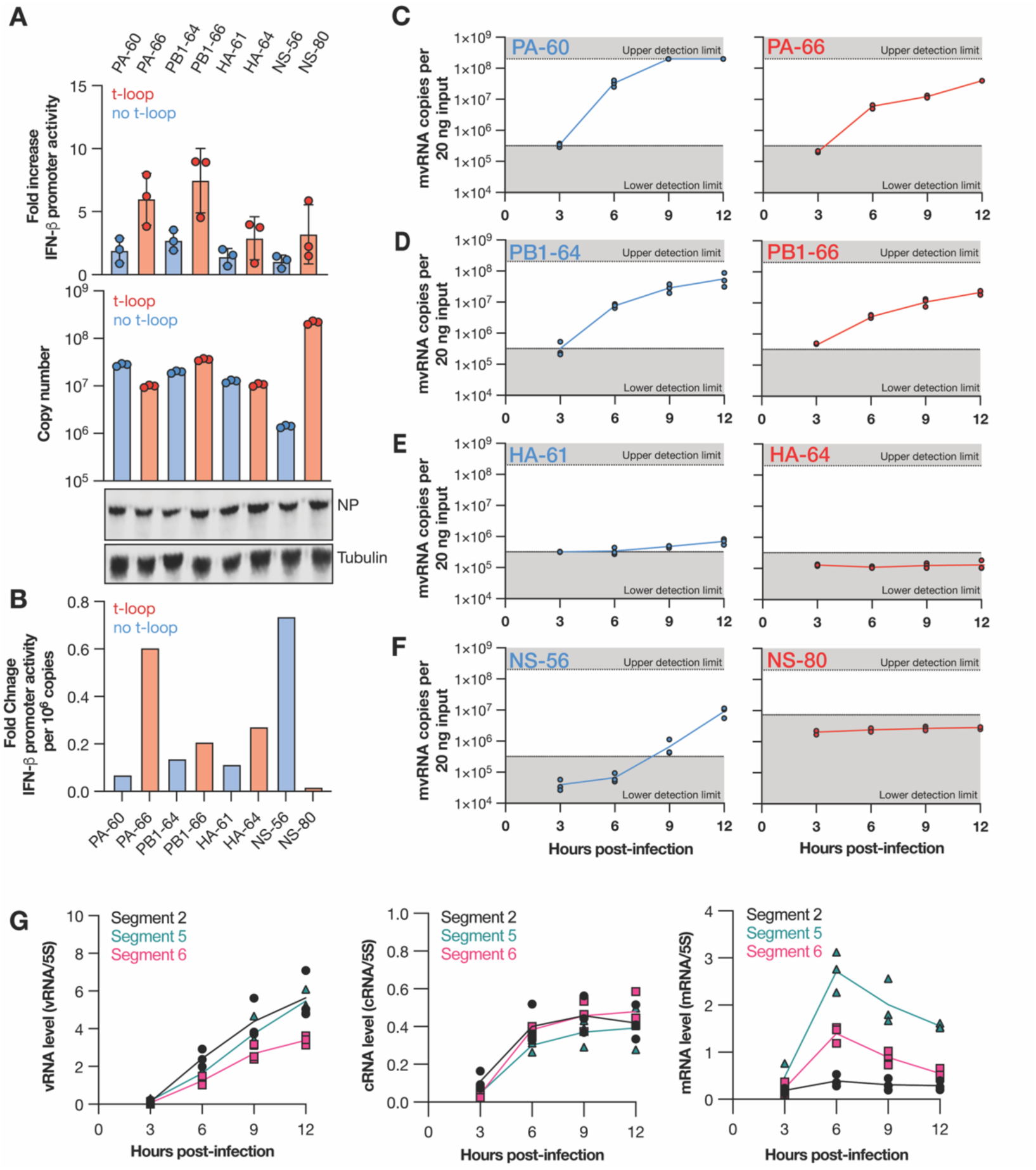
mvRNAs have different effects on IFN-β promoter activity and different kinetics during infection. **A**) IFN-β promoter activation (top graph) and copy number (bottom graph) of transfected mvRNA templates in the presence of the IAV RNA polymerase and NP. Western blot of NP expressed and tubulin loading control in shown in bottom images. mvRNA templates with (red) and without (blue) a t-loop were used. Error bars indicate standard deviation. **B**) IFN-β promoter activation per million mvRNA copies. **C-F**) Copy number of mvRNAs during WSN infection of A549 cells. **G**) Level of replication and transcription products during WSN infection of A549 cells as determined by primer extension. For denaturing PAGE results see Fig. S3. All data represent three biological repeats.

We next designed crRNAs specific for the junction in each mvRNA (Fig. 1B, C) and used these crRNAs together with synthetic RNA standard curves to quantify the number of mvRNA copies present (Fig. 3A). Using the luciferase reporter data and the number of mvRNA copies identified, we were able to calculate the potential of each mvRNA to trigger an innate immune response (Fig. 3B). We observed that per segment, one mvRNA had a strong potential to induce IFNβ promoter activity, while the other mvRNA had a weak potential (Fig. 3B). With one exception (the NS segment), mvRNAs that had a higher copy number tended to have a lower potential to activate the innate immune response (Fig. 3B), which is generally in line with our previous observation on t-loop containing mvRNAs (*9*). The presence of a clear exception on the t-loop model suggests that at least one other, still undiscovered, mechanism must exist through which mvRNAs can activate the innate immune signaling.

Following the transfection-based characterization of the different mvRNAs and confirming our ability to detect them without amplification, we next measured the kinetics of the eight mvRNAs over a synchronized, 12-hour infection of A549 cells with WSN (Fig. 3C-F). We observed that mvRNAs PA-60, PA-66, PB1-64, PB1-66 were detectable in A549 cells as early as 3 hours post infection (hpi) and that their copy number increased steadily over time (Fig. 3C, D). At 9 hpi, the PA-60 signal reached saturation, whereas the PA-66, PB1-64, PB1-66 signals did not reach this level in three biological repeats (Fig. 3D). In contrast, mvRNAs HA-61 and NS-56 were undetectable until 9 hpi (Fig. 3F) and both HA-64 and NS-80 remained undetectable over the course of the 12-hour infection (Fig. 3 E, F). In addition, we observed that three of the mvRNAs with a lower potential to activate the innate immune response (PA-60, PB1-64, and HA-61) were more abundant compared to mvRNAs of the same segment that had a stronger potential to activate the innate immune response (PA-66, PB1-66, and HA-64). Comparing the accumulation kinetics of the PB1 mvRNAs to the full-length segment 2, 5 and 6 kinetics, we found that the mvRNA accumulation followed the trend of the full-length vRNA segments, and not that of the mRNAs of the segments (Fig. 3G, S3).

### PA-60 and PA-66 have different distributions in infected cells

We next sought to compare the subcellular distribution of the PA-60 and PA-66 mvRNA pair over the course of an 8-hour IAV infection. To this end, we performed synchronized infections of A549 cells with WSN and fractionated cells at 2, 3, 4, 5, 6, 7, and 8 hpi (Fig. 4A). During the fractionation process, the volumes were maintained constant to make sure that protein samples could be compared directly to each other (Fig. S4). However, in constant volumes, different fractions have different RNA (background) levels and adsorption levels to the plastics used. To ensure that the LbuCas13a-based reactions for the standard curve (diluted in HEK293T RNA) and the fractionated samples were performed in a similar RNA environment, we normalized the RNA input for each fraction to 75 ng and subsequently analyzed the mvRNA level (Fig. 4A).

**Figure 4.**
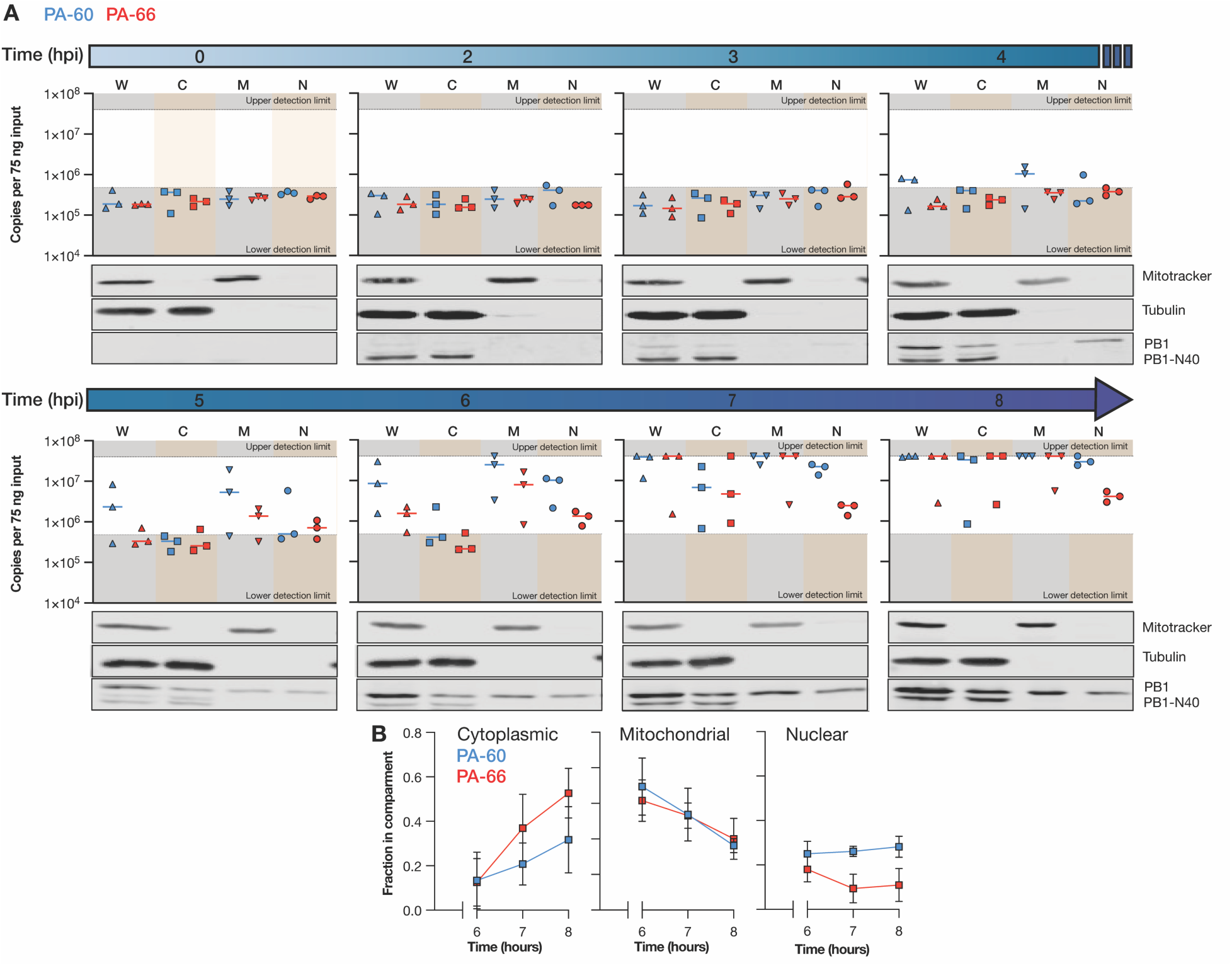
mvRNA PA-60 and PA-66 distribution in subcellular compartments. **A**) Copy number of mvRNAs PA-60 and PA-66 per 75 ng input RNA per subcellular compartment during WSN infection of A549 cells. Data points indicate three biological repeats. Western blots show fractionation of one biological repeat. For additional data see Fig. S4. **B**) Fraction of mvRNAs PA-60 and PA-66 per subcellular compartment. Data points indicate an average of three biological repeats. Error bars indicate SEM. W = whole cell, C = cytoplasmic, M = mitochontrial, N = nuclear.

As shown in Fig. 4A and S4, we were not able to detect mvRNAs before 4 hpi in the whole cell extract, similar to the infection time course described in Fig. 3. From 4 hpi, the PA-60 and PA-66 levels started to increase, with PA-60 showing accumulated more quickly than PA-66. The production of PA-60 and PA-66 followed the detection of PB1 in the nucleus 3 hpi. At 7 hpi, both mvRNAs started to reach the upper limit of our detection assay in nearly all biological repeats (Fig. 4A, B).

In the subcellular fractions (Fig. 4A), we found that PA-60 became first detectable in the nuclear and mitochondrial fractions at 4 hpi. The PA-60 cytoplasmic mvRNA levels remained undetectable until 6 hpi and then rose quickly. At 6 and 7 hpi, the PA-60 nuclear and mitochondrial levels started to the upper detection limit of our assay. The PA-66 mvRNA was slower to accumulate and became first detectable in the nuclear and mitochondrial fractions 5 hpi. At 7 hpi, the PA-66 mvRNA levels became measurable in the cytoplasm and then rose, reaching the upper limit of detection 7 hpi. The PA-66 mitochondrial levels also increased quickly, reaching the upper limit of detection at the same time as the PA-66 cytoplasmic levels. In contrast, the PA-66 nuclear levels deviated from this trend and accumulated only marginally during the course of the infection (Fig. 4A).

To compare the different fractions directly to each other, we corrected the measured LbuCas13a signal for the RNA input adjustment (Fig. S4) and plotted the distribution of the two mvRNAs as a fraction of the total (Fig. 4B). Interestingly, both PA-60 and PA-66 were initially enriched in the mitochondrial fraction. This enrichment then declined for both mvRNAs over the course of the infection. At 8 hpi, PA-60 reached an equal distribution among the three subcellular compartments. However, PA-66 became more enriched in the cytoplasm and only ∼10% of the total signal was found in the nucleus at 8 hpi. This enrichment of PA-66 in the cytoplasm may contribute to the slower accumulation of PA-66 overall, as less PA-66 would be associated with active replication complexes. In addition, the enrichment of PA-66 in the cytoplasm increased its likelihood for detection by cytoplasmic pathogen receptors, such as RIG-I, and thereby its potential for activating the immune response. The enrichment of an immunostimulatory mvRNA in the cytoplasm is also in line with previous data showing that mvRNA templates that activate innate immune signaling become differentially enriched in the cytoplasm over the nucleus (*9*).

### LbuCas13a can detect mvRNAs in RNA extracted from ferret lungs

We had previously shown that mvRNAs can be detected in mouse and ferret lung samples using RT-PCR and next generation sequencing. We next sought to determine whether we could use Cas13 to detect and quantify the highly abundant NP-61 mvRNA in RNA extracted from ferret lung or nasal turbinate tissues infected with pandemic A/Brevig Mission/1/1918 (H1N1). As shown in Fig. 5A, the NP-61 mvRNA could be detected in both tissues on day 1, and in the nasal turbinates on day 3 post infection.

**Figure 5.**
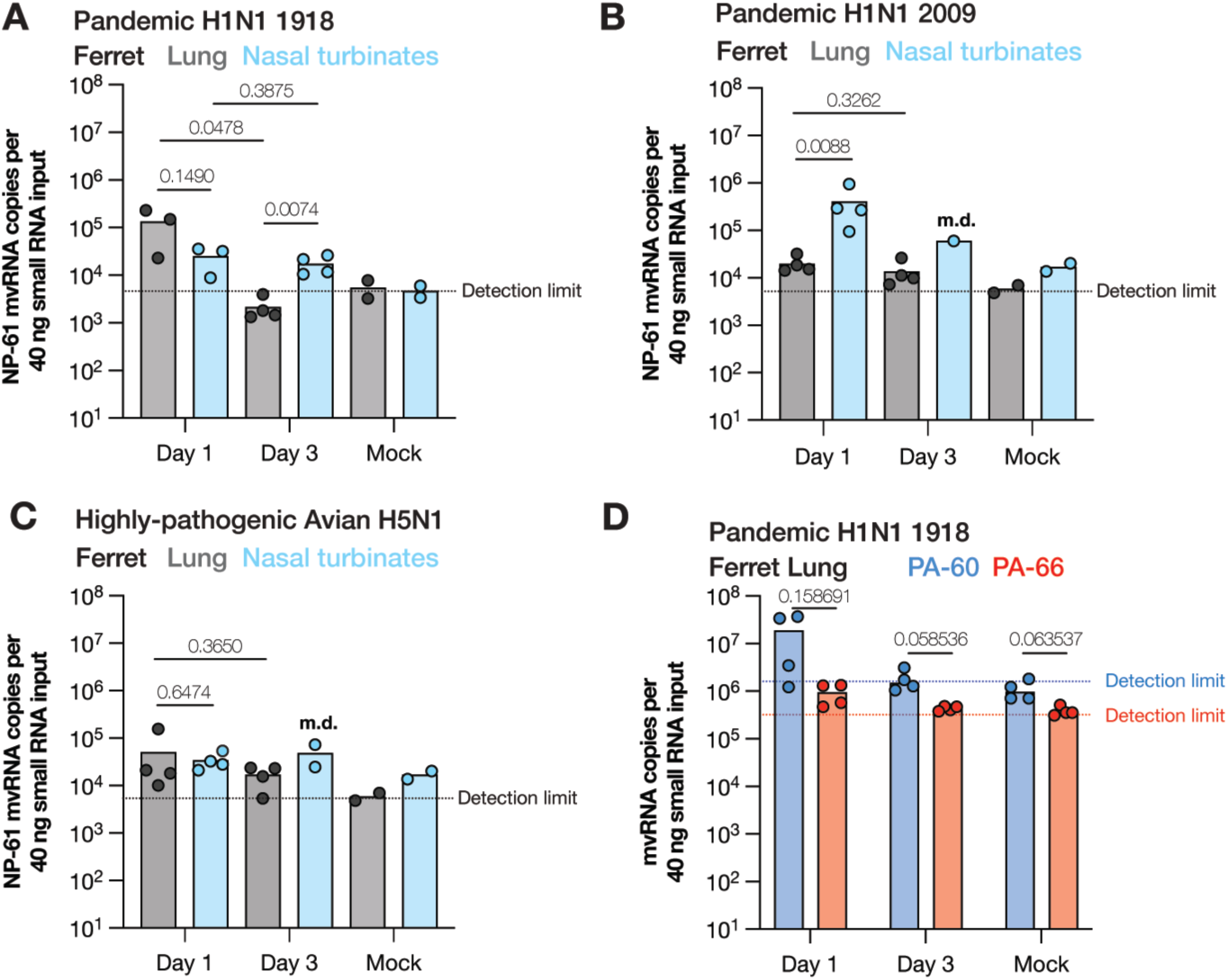
Detection of mvRNAs in ferret lung and nasal turbinates using LbuCas13a. **A**) Detection of mvRNAs NP-61 in small RNA (<200 nt) fractions extracted from ferret lung or nasal turbinates homogenates infected with influenza A/Brevig Mission/1/1918 (H1N1). **B**) Detection of mvRNAs NP-61 in small RNA (<200 nt) fractions extracted from ferret lung or nasal turbinates homogenates infected with influenza A/Indonesia/2005 (H5N1), or **C**) with influenza A/Netherlands/2009 (H1N1). Data points indicate three or four biological infections or mock infections. For ferrets for which not sufficient RNA could not be extracted, missing data points are indicated with m.d. **B**) Detection of mvRNAs PA-60 and PA-66 in small RNA (<200 nt) fractions extracted from ferret lung homogenates infected with influenza A/Brevig Mission/1/1918 (H1N1). Statistical comparisons were performed using One-way ANOVA. P-values are indicated in the panels.

mvRNA sequences vary among IAV strains as the sequences downstream of the promoter are not fully conserved. However, based on our previous deep-sequencing data (*9*), we found that NP-61 was also produced by the pandemic A/Netherlands/602/2009 (H1N1) and highly pathogenic avian A/Indonesia/5/2005 (H5N1) isolates. We therefore next measured the relative abundance of the NP-61 mvRNA in ferret lung and nasal turbinate homogenates infected with these two strains. As shown in Fig. 5B and C, not only were we able to detect NP-61, but we also observed differences in abundance and tissue distribution between the two IAV strains. In line with being an upper respiratory tract virus, the pandemic 2009 IAV strain produced higher NP-61 levels in the nasal turbinates compared to the lungs, whereas avian H5N1 produced equal levels in both parts of the respiratory tract.

We next investigated whether we could also detect PA-60 and PA-66 in the lung tissue, as this is where the impact of innate immune activation is most severe and linked to the cytokine storm. Previous sequence analyses had shown that these two mvRNAs are only produced by H1N1 IAV strains (*9*). As shown in Fig. 5D, both mvRNAs were clearly detectable on day 1 post infection in lungs infected with pandemic A/Brevig Mission/1/1918 (H1N1). Interestingly, the PA-60 mvRNA copy number varied more widely than the more immunogenic PA-66 copy number. On day 3, both mvRNAs had dropped in copy number and were only detectable in some samples. In line with our tissue culture analyses (Fig. 3, 4), we found that the PA-60 copy number was on average higher than the PA-66 mvRNA level (Fig. 5D), but the biological variation among the four ferrets tested was too high for a significant difference.

### LbuCas13a can detect mvRNAs in RNA extracted from clinical swabs

We next investigated whether we could detect mvRNAs in RNA extracted from IAV positive clinical nasopharyngeal swabs. The clinical samples obtained were either positive for seasonal H1N1 or H3N2, and not co-infected with other respiratory pathogens, as determined by clinical RT-qPCR. We first confirmed that we could detect mvRNAs in clinical samples using a RT-PCR and found signals corresponding to the expected size in 6 of 10 IAV positive clinical samples (Fig. S5A). Next, we tested our LbuCas13a assay, using the mvRNA NP-61 mvRNA crRNA. As shown in Fig. S5B and Fig. 6A, we observed positive signals above our limit of detection for 29 of the 30 clinical samples tested. We next used our synthetic NP-61 mvRNA standard curve to calculate the number of NP-61 copies per µl of extracted clinical sample (Fig. 6A). Further analysis showed that the NP-61 copy numbers were not correlated with the amount of RNA input (Fig. 6B), the reported clinical RT-qPCR Ct value (Fig. 6C), or patient age (Fig. 6E). Interestingly, a weak significant difference was observed among the H1N1 samples with respect to the reported patient’s sex (Fig. 6D). Overall, these findings show that mvRNAs can be detected and quantified in a wide range of samples, including influenza virus positive clinical samples.

**Figure 6.**
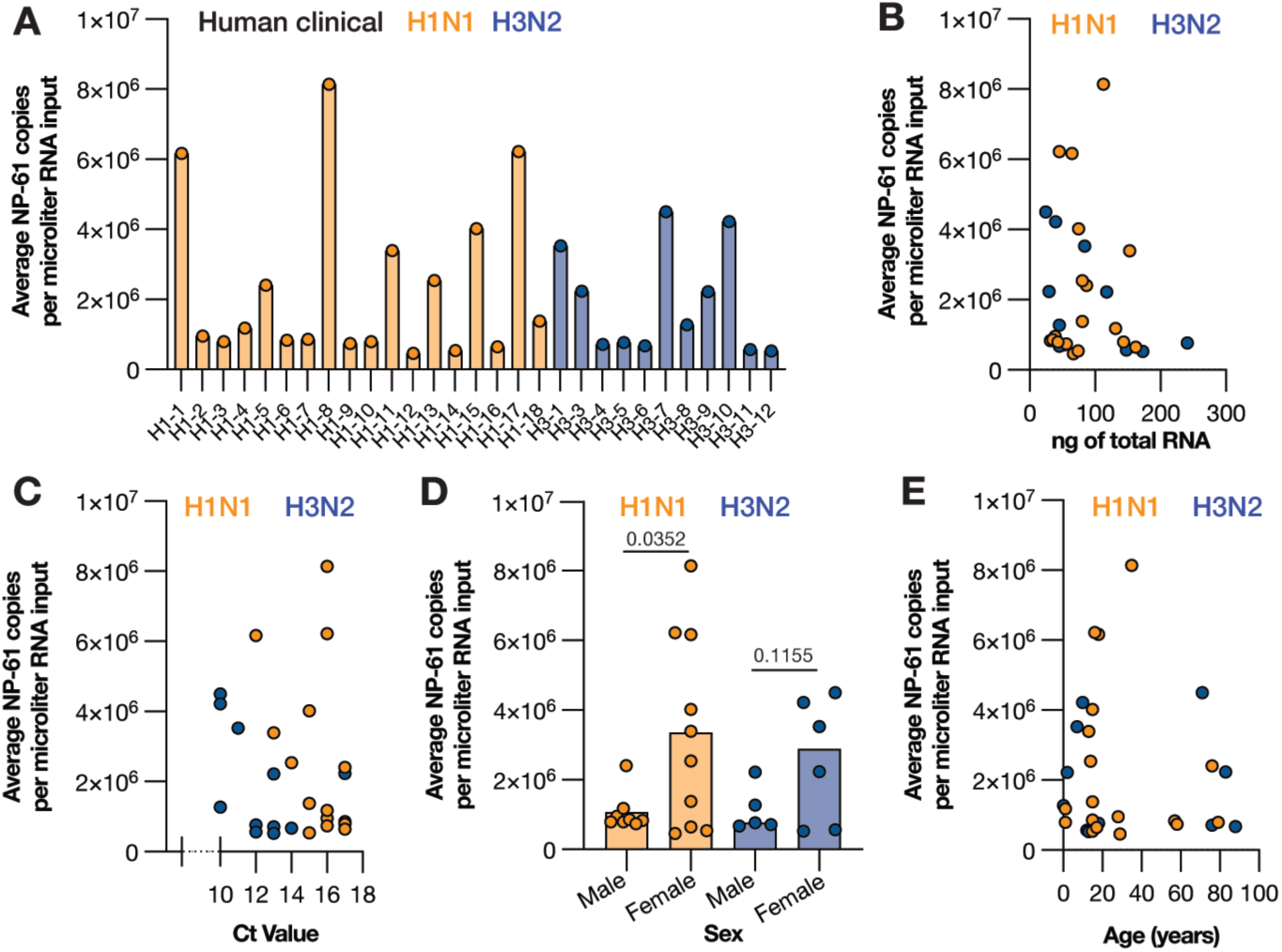
Detection of mvRNAs in clinical nasopharyngeal samples using LbuCas13a. **A**) Copy number of mvRNA NP-61 in equal volumes of total RNA extracted from clinical nasopharyngeal samples for the samples with copy number values above the limit of detection. See table S4 for raw values. **B**) Copy number of mvRNA NP-61 plotted against the amount of ng total RNA used, **C**) plotted against clinical RT-qPCR Ct value, **D**) plotted against the patient sex, and **E**) plotted against patient age. Statistical comparisons in panel D are based on non-parametric t-test.

## Discussion

During IAV infection, aberrant RNA molecules of various lengths and sequences are produced. A subset of these molecules can activate the innate immune response, making them important targets for in-depth characterization. We observed that routinely used RNA quantification methods were sensitive to the mvRNA sequence and failed to faithfully represent mvRNA levels, often creating addition aberrant signals. We therefore explored the RNA-guided enzyme Cas13 as an additional tool to quantify mvRNAs levels in cell culture, animal model, and patient samples.

We first showed that the LbuCas13a ortholog can detect and quantify mvRNAs with a higher sensitivity than primer extension (Fig. 2), and that the LbuCas13a reaction can avoid errors or biases introduced by RT-PCR-based amplification (Fig. 2), because it can detect mvRNAs in an amplification-free manner. In a sample containing a single mvRNA expressed in cells and amplified by the IAV RNA polymerase, LbuCas13 quantifications yielded results that were not significantly different from the primer extension assay (Fig. 2). This result was expected, as any additional aberrant products produced by the IAV RNA polymerase in cell culture (*9*) were excluded in the PAGE-based size selection that underlies the primer extension analysis. Similarly, the TaqMan-based RT-qPCR produced results that were not significantly different from the Cas13-based assay, because the TaqMan probe hybridized to the mvRNA junction similar to the crRNA of Cas13 and thus also excluded any aberrant PCR product that did not include the junction. So, while LbuCas13 is suitable for measuring mvRNA levels with minimal processing and can generate results in a short time with just a plate reader, for samples where RNA levels are low a TaqMan-based RT-qPCR assay may present an alternative. However, when complex mvRNA samples need to be analyzed, which effectively includes all infection experiments, the RT and PCR steps of the TaqMan-based RT-qPCR may introduce unknown, sequence-specific amplification biases that make direct comparisons between mvRNAs in the samples impossible. Using LbuCas13a, we quantified mvRNAs derived from five different IAV segments in transfected cells (Fig. 2 and 3), WSN-infected cells (Fig. 3 and 4), pandemic H1N1 and highly pathogenic H5N1 infected ferret lung and nasal tissues (Fig. 5), and seasonal H1N1 and H3N2 infected clinical samples (Fig. 6). In the tissue culture experiments, we observed that mvRNA dynamics vary among different mvRNA sequences and that they accumulate over the course of infection, similar to the full-length vRNA segments. In agreement with previously published data, we also showed that mvRNAs with a potential to induce a stronger innate immune response are less abundant than mvRNAs that induce a weaker innate immune response for the four segments we examined (Fig. 3). For all but one segment, data also reproduce the finding that mvRNA abundance does not necessarily correlate with innate immune activation and does correlate with the presence of a t-loop (*9*). The exception is the NS segment, whose two mvRNAs do not follow the previous model, suggesting that other mechanisms of innate immune activation by IAV mvRNAs remain to be discovered (Fig. 3). Interestingly, two of the cloned mvRNAs, which had originally been identified through RNA sequencing (*9*), were not detected in A549 cell infections. Presently, it is unclear if these mvRNAs are of exceptionally low abundance and stay below the detection limit, or if these sequences were artifacts of the RT-PCR steps used to make the sequencing library. To address this question and obtain a better estimate of the misidentified mvRNA sequences in the future, a high-throughput, multiplexed version of our assay must be developed and combined with a systematic analysis of all identified (and potentially unknown) mvRNA sequences. These observations are likely also important for studies of DVG sequences, which also rely on RT-PCR-based sequencing libraries to study sequence variations or identify DVG junctions (*8*, *10*).

By examining one immunostimulatory and non-stimulatory mvRNAs derived from segment 3 (PA-66 and PA-60) in more detail, dynamics in their subcellular distribution and accumulation were revealed. We observed that while both stimulatory and non-stimulatory mvRNAs appear in mitochondria early in infection, the immunostimulatory mvRNA then accumulates in the cytoplasm whereas the non-stimulatory mvRNA becomes evenly distributed over the nuclear, cytoplasmic, and mitochondrial fractions (Fig 4B, C). Presently, the mechanism underlying these different distributions is unknown, but the observation agrees with previous analyses of transfected mvRNAs (*9*) and may help explain why the immunostimulatory mvRNA is able to activate the innate immune response better. Whether host or viral factor determine mvRNA export from the nucleus or their distribution across the cell is currently not known. Given the early presence of both mvRNAs in the mitochondrial fraction, it is surprising that we did not observe a strong innate immune response in the presence of both mvRNAs. It is possible that low mvRNA levels are insufficient to activate the innate immune response and that higher mvRNA levels are needed to overcome the negative feedback loops that prevent RIG-I from activating the MAVS-dependent signaling cascade (*28*). Alternatively, mvRNAs associated with the mitochondrial fraction may present species that cannot activate innate immune signaling or that may inhibit innate immune activation through a yet to be discovered process.

Finally, we showed that the difference in abundance between the segment 3 immunostimulatory and non-stimulatory mvRNAs is also present in ferret lungs one day after infection with recombinant 1918 pandemic H1N1 IAV (Fig. 5D). In addition, we were able to measure the copy number of a highly abundant mvRNA in clinical samples that were positive for H1N1 and H3N2 seasonal IAV RNA, demonstrating that our assay works for many different IAV strains and for many different sample types (Fig 6). However, despite detection of this mvRNA in these clinical samples, we are not able to establish any correlations between mvRNA copy number and clinical Ct value, amount of input material, or patient age (Fig. 6). We expect that any correlations were confounded by unknown clinical factors, as well as when patients were tested in the course of their infection, their immune status, sampling differences, and environmental factors. Interestingly, a weak statistical significance was observed between patient sex and mvRNA levels in IAV H1N1 infected patients. Further clinical research will need to be performed to investigate the importance of this difference.

In summary, understanding the dynamics of IAV aberrant RNAs during infection is important because various lines of research have shown that these RNAs can play a role in the outcome of infection. Here we used Cas13 to quantify and characterize 10 different IAV mvRNAs in detail and identify differences in their dynamics and distribution. While our assay is limited in scope due to its limited multiplexing capabilities, it reveals a fascinating, hidden diversity among IAV aberrant RNA molecules, and we hope that our results will stimulate a more in-depth, dynamic characterization of these molecules in order to better understand their role in the outcome of disease.

## Materials and Methods

### Preparation of synthetic RNA and primers

Primers were synthesized by Integrated DNA Technologies (IDT) and resuspended in nuclease-free water to 100 μΜ. Primers were stored at -20 °C and were further diluted prior to analysis. crRNAs were synthesized by IDT and resuspended in nuclease-free water to 100 μM. crRNAs were stored at -70 °C and were further diluted prior to analysis. Synthetic RNA targets were synthesized by IDT and resuspended in nuclease-free water to 100 μM. Synthetic RNA targets were stored at -70 °C and were further diluted prior to analysis.

### Plasmids

The firefly luciferase reporter plasmid under the control of the *IFNβ* promoter [pIFΔ(−116)lucter] and the transfection control plasmids constitutively expressing *Renilla* luciferase under control of the thymidine kinase promoter (pTK-*Renilla*) were described previously (*5*, *9*). The pcDNA3-based WSN protein expression plasmids and pPolI-based WSN template RNA expression plasmids were described previously (*5*, *9*). The mvRNA expressed plasmids were generated by site-directed mutagenesis using the primers listed in Table S3.

### Cells

HEK293T and A549 cells were originally sourced from the American Type Culture Collection (ATCC). All cells were routinely screened for mycoplasma, and grown in Dulbecco’s modified Eagle medium (DMEM) containing pyruvate, high glucose, and L-glutamine (GeneDepot) with 10% fetal bovine serum (Gibco) at 37 °C and 5% CO_2_.

### Transfections

Transfections of HEK293T cells were performed using Lipofectamine 2000 (Invitrogen) and Opti-MEM (Invitrogen) as described previously (*5*, *9*).

### Cell Culture Infections

For infections, 10 cm dishes were seeded with 5 x 10^6^ A549 cells 15 h before infection. Cells were infected with influenza A/WSN/33 at MOI 3. The cells were incubated with virus inoculum at 4°C for 1 h in DMEM/ 0.5% FBS to ensure synchronization of the infections. Next, the inoculum was removed, and the cells incubated at 37 °C in DMEM/ 0.5% FBS for 2, 3, 4, 5, 6, 7, 8 h, or at other times as indicated in the figures. Mock-infected A549 cells were incubated for 8 h. At the relevant time point, cells were lysed in TRI Reagent (Molecular Research Center, Inc) or fractionated using cell Fractionation Kit (Abcam) and subsequently analyzed via RNA extraction.

### RNA Sample preparation

RNAs extraction was carried out using Tri Reagent (Molecular Research Center, Inc) following the manufacturer’s instructions and as described previously (*16*). The RNA concentration was determined using a NanoDrop One spectrophotometer (Thermo Fisher) and diluted in RNase free water prior to analysis.

### RT-qPCR and primer extension

Primers were pre-annealed to the RNA by heating for 2 minutes at 95 °C prior to an RT step carried out using SuperScript III Reverse Transcriptase (Invitrogen) as described previously (*16*). After the RT step, probe-based qPCR was carried out using Luna Universal Probe qPCR Master Mix (New England Biolabs) and the QuantStudio3 Real-Time PCR System (Thermo Fisher) using manufacturer’s instructions. The primers and probes used are listed in Table S3. Primer extensions were performed and analyzed as described previously (*5*, *9*).

### Cas13-based detection reactions

Each reaction contained 10nM LbuCas13a, 4 mM HEPES pH 8.0, 12 mM KCl and 1% PEG, 2U/μL RNAse inhibitor murine (New England Biolabs), 0.25 μM 6UFAM (Table S3), 14mM MgOAc, 5nM crRNA (Table S3), and the reported amount of target RNA. Each reaction was first combined in a volume of 44 μL in 96-well plates. After mixing, 20 μL was transferred in duplicate to 384-well plates. The plate was then placed in a BioTek Cytation 5 Cell Imaging Multi-Mode Reader (Agilent) and incubated at 37 °C for 3 hours. Fluorescence was measured every 5 minutes.

### Curve fitting and Quantification

The maximum slope of each sample and standard was obtained by first determining the slope of every 3 data points starting at t = 0 min (0-10 min, 5-15 min …170 - 180 min). The maximum slopes of the standards were then plotted against the known copy numbers of the standards. Standards that reached saturation or quickly reached saturation (slopehigh-concentration << slopelow-concentration) were omitted prior to fitting the curve. The curve was fit to the Michaelis-Menten equation and K_m_ and V_max_ values were estimated using the Python 3 and the curve-fit trust region reflective (TRF) method from the scipy.optimize package. The V_max_ and K_m_ values were also obtained using Prism 10 software, both Python and Prism 10 software produced the same V_max_ and K_m_ values. Using the obtained K_m_ and V_max_ values and the maximum velocity (V), referred to as the reaction rate in the text, of the sample, the unknown concentrations of the samples were estimated. Limits of detection for each crRNA are listed in Table S2.

### Western blotting and antibodies

IAV proteins were detected using polyclonal anti-IAV PB1 protein antibody (GTX125923, GeneTex) and anti-IAV NP protein antibody (GTX125989, GeneTex), diluted 1:2000 in phosphate buffered saline (PBS)/5% BSA (RPI). To confirm fractionation of the A549 cells, Mitotracker (AB92824, Abcam), γ-tubulin (MCA77G, Bio-rad) and Histone H3 (AB1791, Abcam) antibodies were used, diluted 1:1000-5000 in PBS/5% BSA (RPI). Secondary antibodies IRDye 800 goat anti-rabbit (926-32211, LI-COR), and IRDye 680 goat anti-rat (926-68076, LI-COR) diluted 1:10,000 in PBS/5% BSA were used. Western blots were developed using the Odyssey CLx imaging system (LI-COR).

### Luciferase-based IFN-β promoter activation assays

Transfections to perform RNP reconstitutions and IFN-β reporter assays in HEK293T cells were essentially performed as described previously (*5*, *9*). RNP reconstitutions were carried out in a 6-well plate format with cells (3x10^6^) seeded 24 hours prior to transfection. Per transfection, 250 ng of the pcDNA plasmids encoding PB1, PB2, PA, NP and 250 ng of pPolI plasmid encoding a mvRNA were transfected alongside 100 ng of plasmid expressing firefly luciferase from the IFN-beta promoter and 10 ng of plasmid expressing *Renilla* luciferase using Lipofectamine 2000 (Invitrogen) to the manufacturer’s specifications. Twenty-four hpi, the medium was aspirated, and cells washed with 1 mL of PBS. Cells were resuspended in 200 μl PBS, of which 50 µl was used for the IFN-β promoter activity assay and the remaining cells were used for the Cas13 assay and western analysis. The IFN-β promoter activity assay was done in duplicate using 25 μl of cell suspension in PBS per well in a white 96-well plate format. Next, 25 μl of DualGlo reagent (Promega) was added per well, samples were incubated at RT for 10 min (in dark), and the firefly luciferase readings were taken using a Synergy LX Multimode Microplate Reader (Biotek). Twenty-five μl of Stop-Glo reagent/well was added next, plate was incubated for 10 min at RT (in dark), and the *Renilla* luciferase readings were taken. Firefly luciferase values were normalized by the *Renilla* luciferase values.

### Clinical samples

Nasopharyngeal samples were taken during routine testing from patients hospitalized at Addenbrookes Hospital during the 2016 – 2017, 2017 – 2018, 2018 – 2019 and 2019 – 2020 flu seasons. Patients were positive for either H1N1 or H3N2, but not other respiratory viruses, and samples were taken from a range of pathologies (asymptomatic to death). The study protocol was reviewed and approved by the Health Research Authority (IRAS ID 258438; REC reference 19/EE/0049). Per sample (typically 1.5 mL), 250 µl was used for total RNA extraction using Tri Reagent. RNA was dissolved in 10 µl RNase free water and stored at -70 °C prior to Cas13 analysis.

### Ferret lung samples

Ferret lung samples were kindly provided by Dr. Emmie de Wit and Dr Debby van Riel. Ferret experiments were described previously (*29*) and leftover samples were inactivated and shipped according to standard operating procedures for the removal of specimens from high containment and approved by the Institutional Biosafety Committee. The original experiments were approved by Institutional Animal Care and Use Committee of Rocky Mountain Laboratories, National Institutes of Health, and conducted in an Association for Assessment and Accreditation of Laboratory Animal Care international-accredited facility according to the guidelines and basic principles in the United States Public Health Service Policy on Humane Care and Use of Laboratory Animals, and the Guide for the Care and Use of Laboratory Animals.

### Statistical Testing

GraphPad Prism10 software was used for statistical testing. Unless otherwise stated, error bars represent standard deviation and individual data points indicate biological repeats.

## Acknowledgements

The authors would like to thank members of the Myhrvold and te Velthuis labs for useful discussions and suggestions, Chenyan Huang for preliminary data analysis, and Dr. Emmie de Wit and Dr. Debby van Riel for sharing ferret lung RNA samples.

