## Supplementary material for "Quantification of influenza virus mini viral RNA dynamics using Cas13": Figure S1

### Supplementary Figures

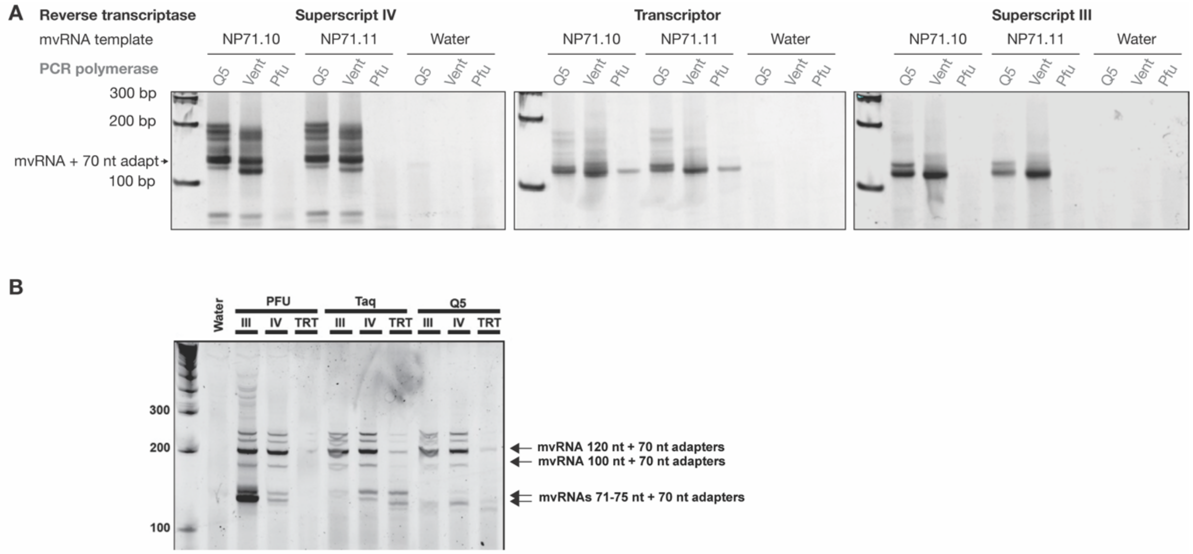


**Figure S1. Detection of synthetic mvRNAs by different RT and PCR enzymes. A**) Two synthetic mvRNAs were reverse transcribed into cDNA by three different RT enzymes and subsequently amplified by three different PCR enzymes. For sequences used, see Table S1. **B**) RT-PCR amplification of 4 different mvRNAs, with lengths 71, 75, 100 and 120 nt, by three different RT and PCR enzymes. Expected sizes and the length of PCR primers are indicated. RT enzymes used were superscript III (III), superscript IV (IV), Transcriptor (TRT). PCR enzymes uses were Pfu polymerase (PFU), Taq polymerase (Taq), and Q5 polymerase (Q5).

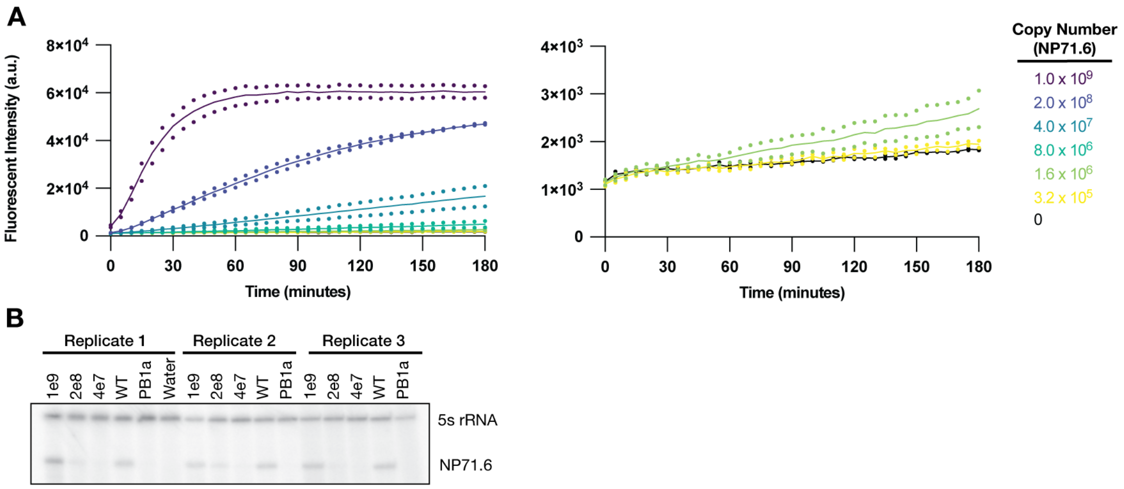


**Figure S2. A**) LbuCas13a detection of synthetic mvRNA NP71.6 diluted in HEK293T total RNA. Right graph is a close-up of bottom part of left graph. **B**) Detection of mvRNA NP71.6 by primer extension following plasmid-based expression in HEK293T cells in the presence of WT or an inactive (PB1a) WSN RNA polymerase. 5S rRNA was used as loading control.

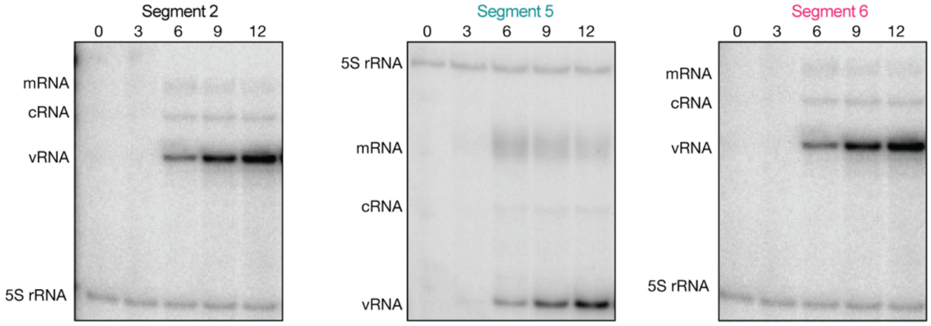


**Figure S3.** Replication and transcription products produced during WSN infection of A549 cells as determined by primer extension and denaturing PAGE. Time point post infection are indicated above each gel image.

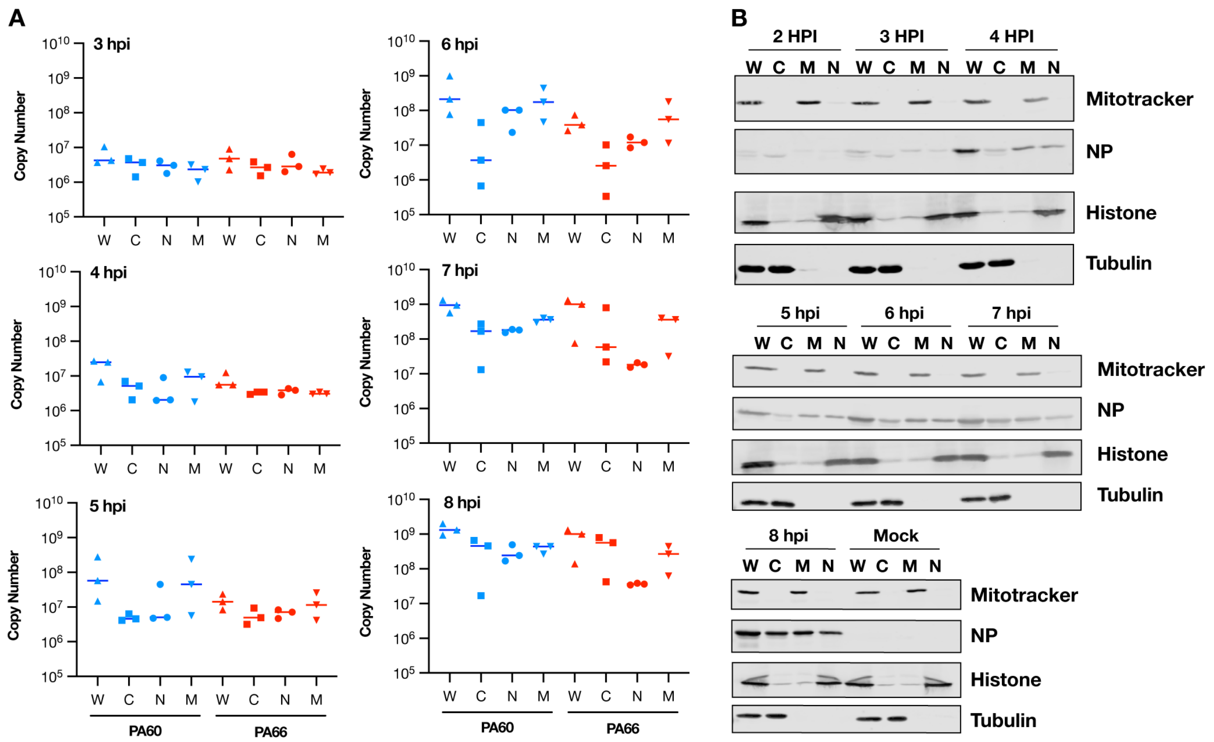


**Figure S4.** Detection of mvRNAs in influenza A virus infection of A549 cells using LbuCas13a. **A**) Cas13 fluorescent signals were converted from copy number per 75 ng of input RNA to the copy number per ul extracted sample to facilitate a direct comparison between the subcellular fractions. **B**) Western blot analysis of WSN infected A549 cell fractionation. Fractions are abbreviated as: W = whole cell, C = cytoplasmic, M = mitochontrial, N = nuclear.

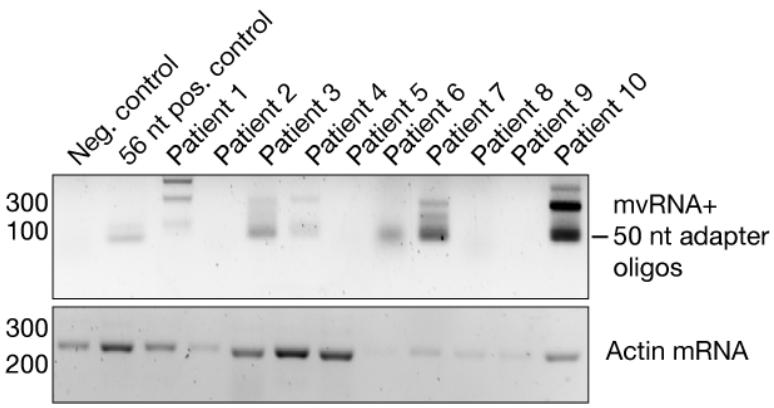


**Figure S5.** Detection of mvRNAs in influenza A virus positive clinical samples using RT-PCR. A 56 nt long segment 5-based mvRNA was used as positive control. As negative control, a clinical sample was used that had tested negative for influenza A virus genetic material.

### Supplementary Tables

**Table S1.** Sequences of mvRNAs studied.

| Name | Lab Reference | Sequence (5′ to 3′) | Figure |
|---|---|---|---|
| NP71.10 | GC50_13 | AGUAGAAACAAGGGUAUUUUUCUUUACUAGUGGCAGCAAAAGCAGGGUAACUAGUCUACCCUGCUUUUGCU | S1A |
| NP71.11 | GC50_15 | AGUAGAAACAAGGGUAUUUUUCUUUACUAGUGGCAGCAAAAGCACCCAUACUAGUCUACCCUGCUUUUGCU | S1A |
| NP71.6 | GC50.9 | AGUAGAAACAAGGGUAUUUUUCUUUACUAGUCCGGCCGUUUUGGUUGCCACUAGUCUACCCUGCUUUUGCU | 2B-2F, S2 |
| NP61 | vNP61 | AGUAGAAACAAGGGUAUUUUUCGAUGUCACUCUGUGAGUGAUUAUCUACCCUGCUUUUGCU | 2G, 5A,5C-D, 6 |
| PA-60 | PA60 | AGUAGAAACAAGGUACUUUUUUGGACAGUAUGCCAUUUUGAAUCAGUACCUGCUUUCGCU | 3A-C, 4, S4, 5B |
| PA-66 | PA66 | AGUAGAAACAAGGUACUUUUUUGGACAGUAUGGAUAGCACAUUUUGAAUCAGUACCUGCUUUCGCU | 3A-C,4, S4, 5B |
| PB1-64 | PB1-64 | AGUAGAAACAAGGCAUUUUUUCAGUCGGAUUGACAUCCAUUCAAAUGGUUUGCCUGCUUUCGCU | 3A-B, D |
| PB1-66 | PB1-66 | AGUAGAAACAAGGCAUUUUUUCAUGAAGGACAAGCUAAACAUUCAAAUGGUUUGCCUGCUUUCGCU | 3A, B, D |
| HA-61 | vHA61 | AGUAGAAACAAGGGUGUUUUUUAGAUCCAUUAUGUCUUUGUCACCCUGCUUUUGCU | 3A, B, E |
| HA-64 | vHA64 | AGUAGAAACAAGGGUGUUUUUUAUCUGAAAGCUUGACACAGUGUUUGGAUCCAUUAUGUCUUUGUCACCCUGCUUUUGCU | 3A, B, E |
| NS-56 | vNS56 | AGUAGAAACAAGGGUGUUUUUCCUUAUAUUUCUGAAAUCCUAAUCUUCCCCUGCUUUUGCU | 3A, B, F |
| NS-80 | vNS80 | AGUAGAAACAAGGGUGUUUUUCCUUAUAUUUCUGAAAUCCUAAUCUCAUUCCCCUGCUUUUGCU | 3A, B, F |
| mvRNA75 | mvRNA75 | AGTAGAAACAAGGGCACTACTGGAAAACTACCTGTTCCATGGCCAACACTTGTCACTACTTTCCCTGCTTTTGCT | S1B |

| | | | |
|---|---|---|---|
| mvRNA100 | mvRNA100 | AGTAGAAACAAGGGTAACAGCTGCTGGGATTACACATGGCATGGATGAACTATACAAATAAATGTCCAGACCTGCAGGCATGCAAGCTCCTGCTTTTGCT | S1B |
| mvRNA120 | mvRNA120 | AGTAGAAACAAGGGCGATGGCCCTGTCCTTTTACCAGACAACCATTACCTGTCCACACAATCTGCCCTTTCGAAAGATCCCAACGAAAAGAGAGACCACATGGTCCTTCCTGCTTTTGCT | S1B |

**Table S2.** crRNA sequences used.

| **Name** | **Target** | **Sequence (5′ to 3′)** | **Limit of Detection (copies)** **(lowest standard that was 3 standard deviations above mock)** | **Figure** |
|---|---|---|---|---|
| CL06 | Human 5s rRNA* | GAUUUAGACUACCCCAAAAACGAAGGGGACUAAAACGGGCGCGUUCAGGGUGGUAUGGCCGUAG | Not determined | 2A |
| CL11 | Human 5s rRNA | GACCACCCCAAAAAUGAAGGGGACUAAAACGGGCGCGUUCAGGGUGGUAUGGCCGUAG | Not determined | 2A |
| CL09 | NP71.6 | GACCACCCCAAAAAUGAAGGGGACUAAAACUAGACUAGUGGCAACCAAAACGGCCGGA | 3e5 | 2B-F, S2A |
| CL51 | NP-61 | GACCACCCCAAAAAUGAAGGGGACUAAAACAGAUAAUCACUCACAGAGUGACAUCGAA | 6e4 | 2G, 5A, C, D, 6 |
| CL12 | PA-60 | GACCACCCCAAAAAUGAAGGGGACUAAAACGGUACUGAUUCAAAAUGGCAUACUGUCC | 3e5 | 3A-C,4, S4, 5A |
| CL13 | PA-66 | GACCACCCCAAAAAUGAAGGGGACUAAAACGGUACUGAUUCAAAAUGUGCUAUCCAUA | 3e5 | 3A-C,4, S4, 5A |
| CL14 | PB1-64 | GACCACCCCAAAAAUGAAGGGGACUAAAACAAUGGAUG | 3e5 | 3A, B, D |

| | | | | |
|---|---|---|---|---|
| | | UCAAUCCGACUGAAAAAAUG | | |
| CL15 | PB1-66 | GACCACCCCAAAAAUGAAGGGGACUAAAACAUUUGAAUGUUUAGCUUGUCCUUCAUGA | 3e5 | 3A, B, D |
| CL25 | HA-61 | GACCACCCCAAAAAUGAAGGGGACUAAAACAAAGCAGGGGAAGAUUAGGAUUUCAGAA | 3e5 | 3A, B, E |
| CL26 | HA-64 | GACCACCCCAAAAAUGAAGGGGACUAAAACAAAGCAGGGGAAUGAGAUUAGGAUUUCA | 3e5 | 3A, B, E |
| CL23 | NS-56 | GACCACCCCAAAAAUGAAGGGGACUAAAACACAAAGACAUAAUGGAUCUAAAAAACAC | 3e5 | 3A, B, F |
| CL24 | NS-80 | GACCACCCCAAAAAUGAAGGGGACUAAAACACUGUGUCAAGCUUUCAGAUAAAAAACA | 8e6 | 3A, B, F |

*LwaCas13a enzyme rather than LbuCas13a enzyme used for the other crRNAs.

**Table S3.** Other oligonucleotides used.

| **Name** | **Description** | **Sequence (5′ to 3′)** | **Figure** |
|---|---|---|---|
| 6UFAM | RNA reporter | /FAM/UUUUUU/3IABKFQ/ | 2, 3, 4, 5, S2, S4 |
| GC- | Targets NP segment | AGCAAAAGCAGGGTAGACTAGT | 2D, 2F, S2B |
| NP 5′ | Targets NP segment | AGTAGAAACAAGGGTATTTTTC | 2D, 2F, S2B |
| GC50.9 probe | Targets NP71.6 | /FAM/TTACTAGTC/ZEN/CGGCCGTTTTGGTTGC/3IABKFQ/ | 2D-F |
| Lv3ga | vRNA RT primer (contains LNA) | GTTCAGACGTGTGCTCTTCCGATCTAGCG+AAAGCAGG | 1A, S1, S5 |
| Lv3aa | vRNA RT primer (contains LNA) | GTTCAGACGTGTGCTCTTCCGATCTAGC+A+AAAGCAGG | 1A, S1, S5 |
| Lv5 | vRNA forward primer (contains LNA) | CACGACGCTCTTCCGATCTHNNNNNNNAGTAGAA+A+CAAGG | 1A, S1, S5 |
| P5short | vDNA fw | GAGATCTACACTCTTTCCCTACACGACGCTCTTCCGATCT | 1A, S1, S5 |
| I7short | vDNA rv | GAGATACTGGTGTGACTGGAGTTCAGACGTGTGCTCTTCCGATCT | 1A, S1, S5 |

**Table S4.** Results of influenza patient clinical sample analysis.

| Fig. Code | Internal Code | Cambridge Code | H1 or H3 | Age (years) | Clinical Ct Value | Sex | Total RNA input (ng) | Copies/ uL |
|---|---|---|---|---|---|---|---|---|
| **H3-1** | C1 | 1193 | H3 | 7 | 11 | F | 83.8 | 3521682.513 |
| **H3-2** | C2 | 1392 | H3 | 62 | 15 | F | 31.5 | Not Detected |
| **H1-1** | C3 | 1109 | H1 | 18 | 12 | F | 64.5 | 6162069.391 |
| **H1-2** | C4 | 1275 | H1 | 28 | 16 | M | 39.1 | 954004.4947 |
| **H3-3** | C5 | 1196 | H3 | 83 | 17 | F | 29.7 | 2238521.406 |
| **H1-3** | C6 | 1163 | H1 | 1 | 17 | M | 143.2 | 786978.3698 |
| **H1-4** | C7 | 1268 | H1 | 1 | 16 | M | 131.2 | 1182889.721 |
| **H3-4** | C8 | 1375 | H3 | 76 | 13 | M | 52.3 | 718879.6206 |
| **H3-5** | C9 | 1497 | H3 | 18 | 12 | F | 241 | 763487.979 |
| **H1-5** | C10 | 1127 | H1 | 76 | 17 | M | 86.7 | 2406890.514 |
| **H3-6** | C11 | 1134 | H3 | 88 | 14 | F | 45.8 | 674301.8301 |
| **H1-6** | C12 | 1135 | H1 | 57 | 17 | F | 31.9 | 838687.1017 |
| **H3-7** | C13 | 1217 | H3 | 71 | 10 | F | 24.4 | 4498742.275 |
| **H3-8** | C14 | 1169 | H3 | 41(days) | 10 | M | 45.3 | 1279877.109 |
| **H1-7** | C15 | 1175 | H1 | 15 | 17 | M | 35.3 | 857500.4898 |
| **H1-8** | C16 | 1139 | H1 | 35 | 16 | F | 113.1 | 8142113.409 |
| **H1-9** | C17 | 1248 | H1 | 58 | 16 | F | 56 | 737658.3474 |
| **H1-10** | C18 | 1219 | H1 | 79 | 17 | M | 43.2 | 794027.1421 |
| **H3-9** | C19 | 1229 | H3 | 2 | 13 | M | 118.1 | 2219306.633 |
| **H1-11** | C20 | 1448 | H1 | 42 | 13 | F | 153.1 | 3399464.279 |
| **H1-12** | C21 | 1126 | H1 | 84 | 29 | F | 67.0 | Not detected |
| **H3-10** | C22 | 1588 | H3 | 82 | 10 | F | 39.9 | 4215913.226 |
| **H1-13** | C23 | 1464 | H1 | 87 | 14 | F | 80.6 | 2535752.682 |
| **H1-14** | C24 | 1403 | H1 | 34 | 15 | F | 74.1 | 545581.3123 |
| **H3-11** | C25 | 1608 | H3 | 87 | 12 | F | 148.0 | 564835.1674 |
| **H1-15** | C26 | 1633 | H1 | 29 | 15 | F | 75.0 | 4014859.47 |
| **H3-12** | C27 | 1673 | H3 | 32 | 13 | F | 173.1 | 531682.0197 |
| **H1-16** | C28 | 1314 | H1 | 23 | 17 | F | 162.2 | 640879.3779 |
| **H1-17** | C29 | 1435 | H1 | 3 | 16 | F | 45.2 | 6224905.957 |
| **H1-18** | C30 | 1689 | H1 | 81 | 15 | F | 80.3 | 1378330.248 |
